# Genome-wide association studies reveal distinct genetic correlates and increased heritability of antimicrobial resistance in *Vibrio cholerae* under anaerobic conditions

**DOI:** 10.1101/2021.06.26.450051

**Authors:** A. Creasy-Marrazzo, M.M. Saber, M. Kamat, L. S. Bailey, L. Brinkley, E. T. Cato, Y. Begum, M.M. Rashid, A. I. Khan, F. Qadri, K. B. Basso, B. J. Shapiro, E. J. Nelson

**Affiliations:** Departments of Pediatrics and Environmental and Global Health, University of Florida, Gainesville, FL, USA; Department of Microbiology & Immunology, McGill University; Departments of Chemistry, University of Florida, Gainesville, FL, USA; Infectious Diseases Division (IDD) & Nutrition and Clinical Services Division (NCSD), International Centre for Diarrhoeal Disease Research, Bangladesh (icddr,b), Dhaka, Bangladesh

**Keywords:** Antimicrobial resistance, AMR, antibiotics, cholera, diarrhoea, diarrhea, *Vibrio cholerae*, enteropathogens, Bangladesh, anaerobic, anoxic, hypoxic, respiration

## Abstract

The antibiotic formulary is threatened by high rates of antimicrobial resistance (AMR) among enteropathogens. Enteric bacteria are exposed to anaerobic conditions within the gastrointestinal tract, yet little is known about how oxygen exposure influences AMR. The facultative anaerobe *Vibrio cholerae* was chosen as a model to address this knowledge gap. We obtained *V. cholerae* isolates from 66 cholera patients, sequenced their genomes, and grew them under anaerobic and aerobic conditions with and without three clinically relevant antibiotics (ciprofloxacin, azithromycin, doxycycline). For ciprofloxacin and azithromycin, the minimal inhibitory concentration (MIC) increased under anaerobic conditions compared to aerobic conditions. Using standard resistance breakpoints, the odds of classifying isolates as resistant increased over 10 times for ciprofloxacin and 100 times for azithromycin under anaerobic conditions compared to aerobic conditions. For doxycycline, nearly all isolates were sensitive under both conditions. Using genome-wide association studies (GWAS), we found associations between genetic elements and AMR phenotypes that varied by oxygen exposure and antibiotic concentrations. These AMR phenotypes were more heritable, and the AMR-associated genetic elements were more often discovered, under anaerobic conditions. These AMR-associated genetic elements are promising targets for future mechanistic research. Our findings provide a rationale to determine if increased MICs under anaerobic conditions are associated with therapeutic failures and/or microbial escape in cholera patients. If so, there may be a need to determine new AMR breakpoints for anaerobic conditions.

**Impact statement:** Many bacterial pathogens experience anaerobic conditions in the gut, but antimicrobial resistance (AMR) phenotypes are generally tested under ambient aerobic conditions in the laboratory. To better understand AMR under conditions more similar to natural infections, we used *Vibrio cholerae* as a model enteric pathogen. We sequenced the genomes and assessed the growth of *V. cholerae* isolates with different concentrations of three antibiotics, under anaerobic and aerobic conditions. In support of the hypothesis that AMR varies according to oxygen exposure, *V. cholerae* was more resistant to antibiotics under anaerobic conditions. We found many previously known genes associated with resistance; however, some of these genes were only resistance-associated under aerobic conditions. Resistance to azithromycin and doxycycline only had a detectable genetic component under anaerobic conditions. Together, our results point to distinct genetic mechanisms of resistance under anaerobic conditions and suggest several candidate genes for experimental follow-up.

**Data summary:** All sequencing data generated in this study are available in NCBI under BioProject PRJNA818081.

## INTRODUCTION

Clinically relevant laboratory methods are essential to gauge the extent to which the antibiotic formulary is threatened by antimicrobial resistance (AMR). Knowledge gaps remain on the degree to which *in vitro* AMR assays reflect *in vivo* AMR physiology. Facultative anaerobic pathogens experience hypoxia and anoxia within the gastrointestinal tract, yet AMR assays rely on aerobic conditions [1]. How oxygen exposure effects AMR is poorly understood. To investigate this question, we chose the facultative anaerobe *Vibrio cholerae* as a model system. In *V. cholerae*, classic mechanisms for AMR, and physiologic pathways for anaerobic respiration and fermentation, are well characterized [2-9]. The disease cholera is also one of the few non-invasive diarrheal diseases for which antibiotics are indicated, albeit conditionally [10-12].

Rehydration is the definitive intervention for acute diarrheal disease [11]; antibiotics are supportive and indicated for only a few diarrheal diseases, including cholera. The World Health Organization (WHO) recommends ciprofloxacin, azithromycin or doxycycline for cholera patients with severe dehydration [10-12]; antibiotics shorten the frequency and duration of diarrhea. In practice, guideline adherence in cholera endemic regions may be low out of clinical concern that a patient ‘might’ have cholera and may develop severe dehydration, contributing to rates of inappropriate antibiotic usage that can rise above 90% [13, 14]. Strong regional associations between antibiotic use and rise of AMR have been observed across enteric taxa [15, 16]. Given that AMR genes frequently co-localize on mobile elements [17], inappropriate single-agent therapy poses a risk of multidrug-resistance (MDR) selection.

Associations between AMR phenotypes and genotypes are known for the three antibiotics recommended to treat cholera; the cognate AMR mechanisms share commonality across Gram negative taxa. Ciprofloxacin (a fluoroquinolone) resistance mechanisms include mutations in genes encoding type II topoisomerases: heterotetrameric DNA gyrase (GyrA_2_GyrB_2_) and DNA topoisomerase IV (ParC_2_ParE_2_). Mutations in the quinolone resistance-determining region (QRDR) of *gyrA* and *parC* can yield additive resistance phenotypes [18]. Fluoroquinolone resistance can also arise by efflux pump upregulation, by downregulation of outer membrane porins that permit quinolone entry, and by the expression of the quinolone resistance protein (Qnr, a pentapeptide repeat protein) that protects the target gyrase protein [18]. Resistance can increase over 30-fold compared to wild-type when strains harbor *qnr* family genes.

In *V. cholerae*, diverse AMR genes, including *qnr*, often reside on an integrative and conjugative element (ICE; ‘SXT’ in *V. cholerae*) [17, 19, 20]. Azithromycin (a macrolide) resistance mechanisms are similarly diverse and include mutations in the 23S ribosomal RNA (rRNA) target genes and ribosomal protein genes. Macrolide resistance is conveyed by carriage of rRNA methyltransferase genes (*erm*) and associated induction mechanisms, *cis-*acting peptides, efflux systems (e.g. *mef, msr)*, macrolide esterases (e.g. *ere*), and macrolide phosphotransferases such as *mphA* which can reside on the *V. cholerae* SXT element [21]. Doxycycline (a tetracycline) resistance is conferred by mutations in the 16S rRNA component of the 30S ribosomal subunit [22]. Additional mechanisms include tetracycline-specific ribosomal protection proteins (RPPs), tetracycline specific efflux pumps (e.g. *tet*(59)) which can reside on SXT element intrinsic efflux pumps, AraC-family transcriptional activators (e.g. MarA), and cytoplasmic ATP-dependent serine proteases [22].

Associations between AMR phenotypes and genotypes have been studied by random mutagenesis, phenotypic screening, and network analyses [23-28], and applied in *V. cholerae* [29]. These approaches uncover how the effect of an antibiotic is shaped by a large number of often more subtle physiologic perturbations, including altered DNA synthesis/repair, central metabolism/growth, and SOS response [30, 31]. AMR assays conducted under aerobic conditions alone may not reflect these physiologic perturbations experienced in the host. Within bacteria, aerobic oxidative phosphorylation generates reactive oxygen species (ROS) that are lethal unless a sufficient defense is mounted by factors like superoxide dismutase, catalase, and glutathione systems [32, 33]. Under anaerobic conditions, growth rate typically slows and proton motive force is reduced [34, 35], which can have both synergistic and antagonistic effects on antibiotics [31, 36]. In *Escherichia coli*, ROS are generated after fluroquinolone treatment under aerobic conditions [37] and fluoroquinolone resistance increases under anaerobic conditions [38, 39]. The extent to which tetracyclines and macrolides induce ROS and how anaerobiosis influences resistance and susceptibility is less known [30, 40].

The objective of this study was to compare AMR phenotypes, with underlying genotypes, under aerobic and anaerobic conditions among isolates obtained from cholera patients. The study rationale assumes that the lower gastrointestinal tract of cholera patients is hypoxic/anaerobic, despite animal experiments that suggest aerobic respiration in the upper gastrointestinal tract is important for infection [41, 42]. Using minimal inhibitory concentration assays (MICs), we found that AMR increased under anaerobic conditions for select antibiotics, and novel genetic targets for AMR were discovered under anaerobic conditions.

## METHODS

### Clinical sample collection

The two sample collections analyzed were part of previously published IRB approved studies [13, 43]. In the primary collection, stool samples were obtained during the spring cholera outbreak period of 2006 at the International Centre for Diarrhoeal Disease Research, Bangladesh (icddr,b) in Dhaka, Bangladesh. Samples were collected prior to hospital administration of antibiotics; patient histories were negative for known antibiotic exposure. The library consisted of 67 *V. cholerae* isolates (Supplementary Table 1); paired stool supernatant for mass spectrometry was available for 50 isolates. In the secondary collection, samples were obtained in 2018 as part of a cholera surveillance study conducted across Bangladesh [13]. Samples were collected at hospital admission independent of reported antibiotic exposure; 277 out of 282 isolates cultured and were analyzed to assess generalizability of the AMR profiles identified in the primary collection.

### Antimicrobial resistance testing

Growth kinetics and the MIC determinations for ciprofloxacin, azithromycin, and doxycycline were performed on isolates from the primary collection in LB broth with twelve two-fold serial dilutions with concentrations spanning the CLSI MIC breakpoints [1] for *V. cholerae* (ciprofloxacin = 2 μg/ml; azithromycin = 8 μg/ml; doxycycline = 8 μg/ml) [1]. Isolates were prepared and grown aerobically at 37°C in 15-ml tubes containing 5-ml LB broth at 220 rpm. Bacteria were back-diluted to a final optical density (OD) 600 nm of 0.01 (200 μl/well) in LB with or without the respective antibiotic dilution-series in black Corning CoStar clear-bottom 96-well plates. Plates were placed in a BioTek Synergy H1 reader pre-warmed to 37°C with the lid on. Anaerobic conditions were generated using a continuous chamber flow (5% CO_2_, 95% N_2_) and a BioTek CO_2_/O_2_ gas controller; anaerobic growth plates were given a 10-minute equilibration period. OD 600 nm was measured every 2 minutes for 8 hours at 37°C with orbital shaking at 220 rpm. A standard logistic equation was fit to growth curve data using the R package *growthcurver* version 0.3.0 [44]. Outcome measures were intrinsic growth velocity (growth rate that would occur if there were no restrictions on total population size), carrying capacity (K; maximum possible population size), and area under the curve (AUC). The MIC was determined using a logistic fit for growth over the twelve, two-fold serial dilutions of the test antibiotic. Binary phenotypic sensitive/resistance categories were set in concordance with Clinical and Laboratory Standards Institute (CLSI; M45 2018 3^rd^ edition) [45]. In general, CLSI breakpoints are set by clinical and bacteriological response data, pharmacokinetic and pharmacodynamic simulations, and expert working group experience. The breakpoints for *V. cholerae* are based on aerobic assays. Susceptible (sensitive) is defined by CLSI as the “category that implies that isolates are inhibited by the usually achievable concentrations of antimicrobial agent when the dosage recommended to treat the site of infection is used”. Those not sensitive were scored as resistant (combines indeterminate and resistant). Three isolates and one reference strain (E7946) were used to assess media acidification during anaerobic and aerobic growth; the pH was measured using dipsticks (pH range 5.2-7.2). The assays were conducted with and without the addition of 20mM fumarate as an alternative electron acceptor for anaerobic respiration.

### Use of catalase to determine if ROS contribute to antibiotic sensitivity

To test if the reduction of ROS was associated with increased resistance to antibiotics under aerobic conditions, MICs were determined for two select strains (E7946, EN160) with and without catalase (10 U/ml; final concentration) added to the media. Growth curves were performed with viable counts as endpoints to determine the minimum dose for lethality by H_2_O_2_ or protection by catalase.

### Whole-genome sequencing

Genomic DNA was extracted from *V. cholerae* isolates from the primary collection using the Qiagen DNeasy Blood and Tissue Kit. Library construction was completed using the Illumina Nextera XT v.2 DNA Library Preparation Kit. Libraries were sequenced in three illumina MiSeq runs. Two batches of twenty-four genomes and one batch of nineteen were pooled and sequenced on a MiSeq for 500 cycles per run. Using CLC Genomics Workbench v20, raw reads were filtered by length, trimmed, and mapped to the reference genome (*V. cholerae* O1 El Tor E7946) to identify single-nucleotide variants. Of the 67 isolates, 66 yielded sufficient coverage (>50X) of the *V. cholerae* genome. We proceeded with these 66 genomes for further analysis. To identify genes not present in the reference genome, contigs were assembled *de novo* using CLC Genomics Workbench v20.

### Genome-wide association studies (GWAS)

To extract genomic variants capturing all sources of variation in the genome (i.e. single nucleotide variants, indels and gene presence/absence) without *a priori* assumption about the underlying gene content of each sample (e.g. accessory genes or plasmids), unitigs were generated from the 66 genomes assembled using GATB [46]. Unitigs are sequences of variable length (unlike k-mers of fixed length *k*) which represent the variations in the population of genomes under study in high-resolution. GWAS were performed using linear mixed models implemented in pyseer v.1.3.6 and adjusted for population stratification using the kinship matrix estimated from the phylogenetic tree[47].

To generate the phylogenetic tree, genome alignments consisting entirely of variable nucleotides were produced from whole genome SNP data generated by CLC Genomics Workbench v20 using VCF-kit 0.1.6 [48]. The tree was then inferred by RaxML under the general time reversible (GTR) model with rate variation across sites following a GAMMA distribution[49]. We used the linear-mixed model approach to adjust for population stratification and linkage disequilibrium in microbial GWAS [50]. Heritability (*h*^*2*^), an estimate of the proportion of the phenotype variance that can be explained by total genomic variation represented in the unitigs, was also calculated using pyseer v.1.3.6. Likelihood-ratio test *p*-values for the association tests were adjusted for multiple-testing by Bonferroni correction (at a genome-wide false discovery rate of 0.05) for the number of unique unitig patterns (i.e. only giving one count to a unitig with an identical presence/absence profile across genomes). We also removed unitigs tagged with the errors ‘bad-chisq’, ‘pre-filtering-failed’, ‘lrt-filtering-failed’, ‘firth-fail’ and ‘matrix-inversionerror’ after the analysis. To further remove false positive GWAS hits, we removed any considerable clusters of unitigs (> 20) with identical *p-*values, as these are likely to be lineage-specific markers or markers with strong linkage disequilibrium comprised of mostly non-causal variants linked on the same clonal frame. GWAS hits were annotated by mapping the unitigs to two reference genomes of *V. cholerae*, namely, E7946 (NCBI assembly accession number: GCA_002749635.1) and O1 biotype El Tor strain N16961 (NCBI assembly accession number: GCA_003063785.1) using BWA. Statistically significant GWAS hits were further annotated with the CARD resistance gene identifier (RGI) after filtering the ‘loose’ hits and hits with identity <0.90.

### Antibiotic detection by liquid chromatography mass spectrometry (LC-MS/MS)

The approach was based on a prior study [51]. Stool supernatant from the primary collection were obtained by centrifugation and filtration (0.2 μM surfactant-free cellulose acetate; Thermo Scientific Nalgene). Proteins were precipitated (1:7 ratio (v/v) of water:methanol). Supernatants were diluted with methanol and water (1:1 v/v) in 0.1% formic acid for liquid chromatography, and 5 µl of supernatant was injected for analysis. LC/MSMS was performed on a 2.1 × 150-mm Hypersil Gold aQ column (particle size, 3 μm) using a high-performance liquid chromatography system (Thermo UltiMate 3000 series) with an LTQ XL ion trap mass spectrometer (Thermo Fisher Scientific). Mobile phases were 1% formic acid in water (A) and 1% formic acid in methanol (B) and held at a constant 5%B for 2min before ramping to 95%B at 15 min where it was held for an additional minute before returning to starting conditions for a total run time of 25 min.

Eluent was ionized using electrospray ionization (ESI) in positive mode at a spray voltage of 5 kV, a nitrogen sheath gas flow rate of 8 L min^-1^, and capillary temperature of 300°C. Two scan events were programmed to perform an initial scan from *m/z* 100 to 1000, which was followed by targeted collision induced dissociation based on a retention time and mass list. Retention time windows ranged from 0.35 minutes to 6.50 min, depending on the elution range of the standards at high and low concentrations.

Masses were targeted for the most abundant adduct or ion associated with each antibiotic (typically the [M+H]^+^ ion) with a *m/z* 1 window. Data analysis for amoxicillin, sulfamethoxazole/trimethoprim, azithromycin, tetracycline, doxycycline, metronidazole, nalidixic acid, and ciprofloxacin was performed manually using extracted ion chromatograms and MSMS matching with an in-house antibiotic MSMS library using Xcalibur 2.2 SP 1.48 (Thermo Fisher Scientific).

### Statistical analysis

Bivariate analyses of categorical data were analyzed using Fisher’s Exact Test, and continuous data were analyzed using the Mann-Whitney U Test (alpha = 0.05). McNemar’s test was used to analyze paired data (alpha = 0.05).

## RESULTS

### Comparison of antimicrobial resistance under aerobic and anaerobic conditions

We measured baseline growth parameters for each isolate under anaerobic and aerobic conditions and found that carrying capacity, area under the growth curve (AUC), and growth velocity were all significantly lower under anaerobic conditions (Supplementary Table 2). We tested an assumption that under anaerobic conditions mixed fermentation and anaerobic respiration would occur in LB media. Analyzing a subset of three isolates and the reference strain E7946, we monitored for acidification as a sign of fermentation and assessed the impact of the addition of an alternative electron acceptor (20 mM fumarate) for anaerobic respiration. Under anaerobic conditions, one out of four strains acidified the LB media to 6.0 (Supplementary Table 3), suggesting fermentation in at least some isolates. The addition of fumarate as an alternative electron acceptor under anaerobic conditions resulted in a small increase in AUC of 25-27% (Supplementary Table 3), suggesting that in LB media anaerobic respiration likely occurs but is limited for alternative electron acceptors. These results are consistent with mixed fermentation and anerobic respiration in our experimental conditions.

In this physiologic context and using standard antibiotic breakpoints established for aerobic conditions, AMR differed between anaerobic versus aerobic conditions (Fig. 1); distributions of single and multi-agent AMR phenotypes are shown (Fig. 2A). The MIC modes for ciprofloxacin were 8 μg/ml (min=0.016 μg/ml; max=32 μg/ml) and 2 μg/ml (min=0.004 μg/ml; max=8 μg/ml) under anaerobic and aerobic conditions, respectively (Supplementary Table 4); the rates of resistance under anaerobic (93%; N=62/67) and aerobic (54%; N=36/67) conditions were significantly different (Fig. 1A; McNemar’s test p<0.001; Supplementary Table 5). For azithromycin, the MIC modes were 32 μg/ml (min= 8 μg/ml; max=124 μg/ml) and 4 μg/ml (min=1 μg/ml; max=32 μg/ml) under anaerobic and aerobic conditions respectively (Supplementary Table 4). The rates of resistance under anaerobic (n=67/67; 100%) and aerobic (n=15/67; 22%) conditions were significantly different (Fig. 1B; McNemar’s test p<0.001; Supplementary Table 5). For doxycycline, the MIC modes were 1 μg/ml under both aerobic and anaerobic conditions, respectively (Supplementary Table 4); one isolate was resistant under anaerobic conditions alone, and one isolate was resistant under both anaerobic and aerobic conditions. The odds of classifying isolates as resistant increased over 10 times for ciprofloxacin (OR= 10.5; 95% CI= 3.61-37.7) and over 200 times for azithromycin (OR = 213; 95% CI= 31.9->5000) under anaerobic compared to aerobic conditions.

**FIG 1.**
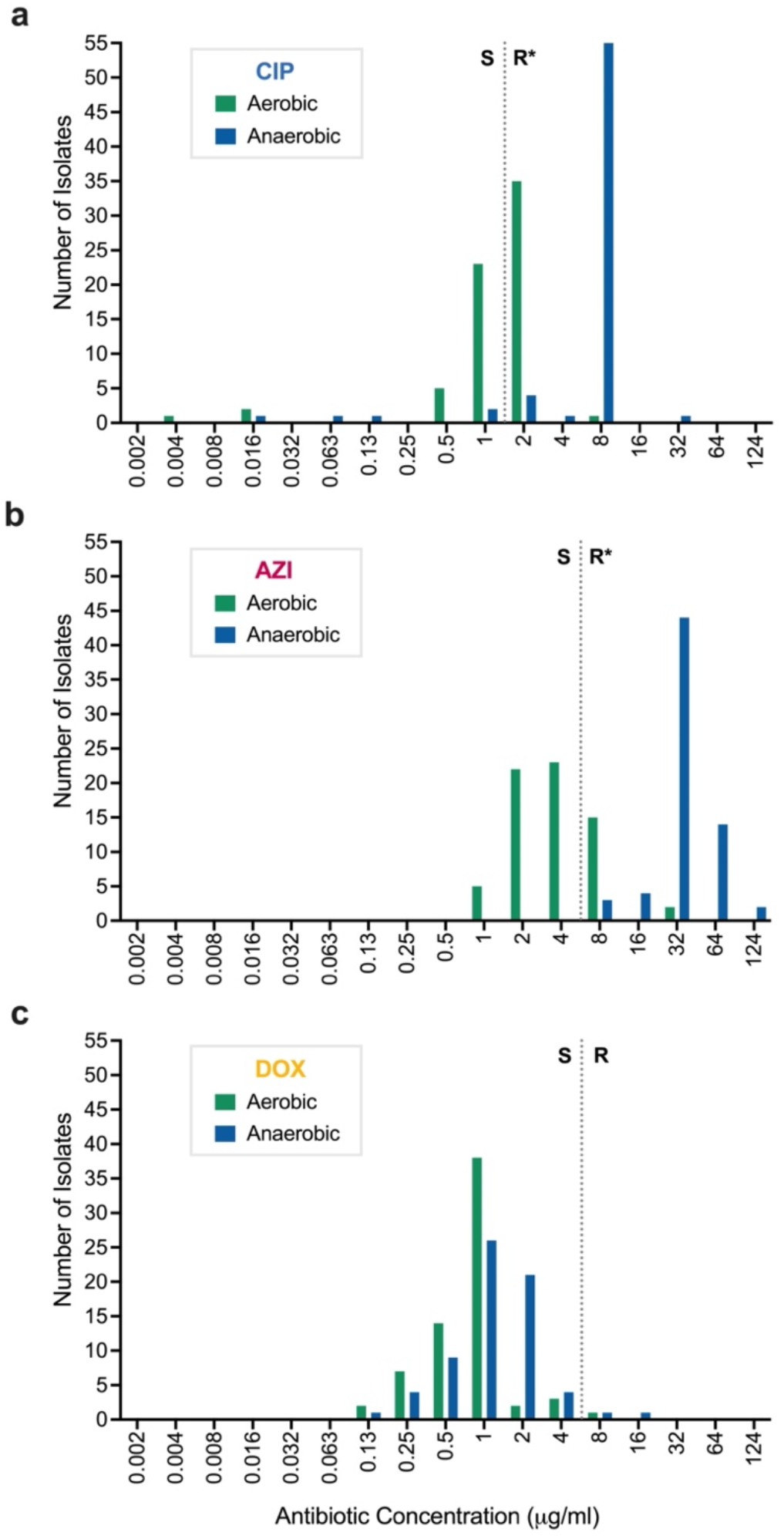
Distribution of minimal inhibitory concentrations (MICs) under aerobic and anaerobic conditions among clinical isolates from the primary collection. Ciprofloxacin (CIP; **a**), Azithromycin (AZI; **b**), and Doxycycline (DOX; **c**). Data are from 67 human-shed *V. cholerae* isolates. The MICs for each isolate under aerobic (green) and anaerobic (blue) conditions were enumerated, and the number of isolates with a given MIC (µg/ml) are represented as bars. Dotted lines are the breakpoint for resistance per CLSI standards which are based on assays under aerobic conditions (CIP = 2 µg/ml; AZI = 8 µg/ml; DOX = 8 µg/ml). S=sensitive. R = Resistant. “*” represents a significant difference in the frequency of isolates identified as resistant to ciprofloxacin and azithromycin by McNemar’s test (both p<0.001).

**FIG 2.**
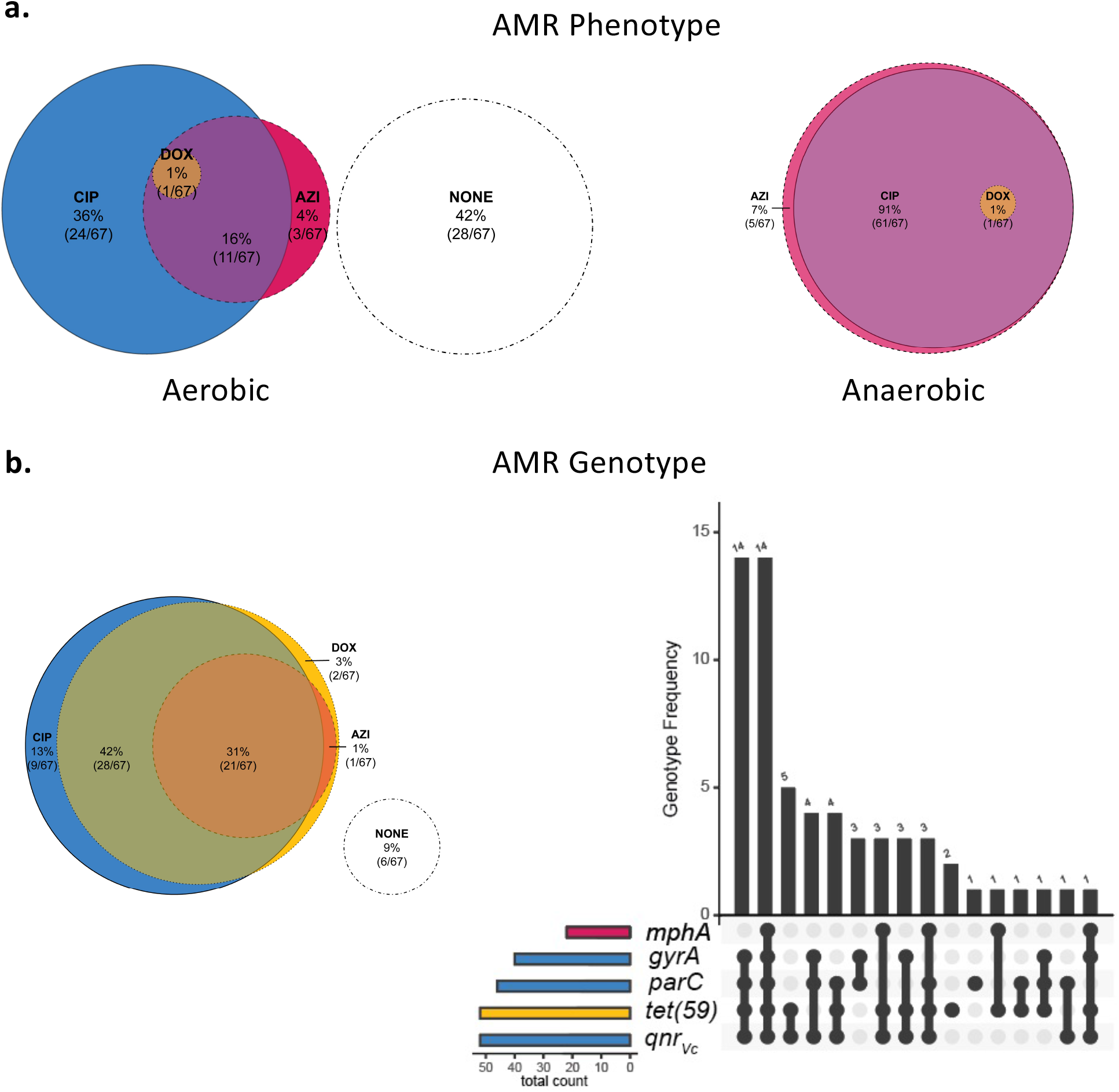
AMR phenotypes and known AMR genetic elements in human-shed *V. cholerae* isolates from the primary collection. Pink, yellow, and blue color coding is used to respectively indicate AMR phenotypes and genotypes to azithromycin (AZI), doxycycline (DOX) and ciprofloxacin (CIP); colors blend when overlapped. Isolates with no resistance (sensitive) are shown in white circles. (**a**) Proportional Venn diagram (Euler) of AMR phenotypes to AZI, DOX, and/or CIP under aerobic (left) and anaerobic conditions (right). Counts indicate the number of isolates with the corresponding phenotype. (**b**) AMR genotypes from known resistance genes in whole genome sequences. Shown is the distribution of known AMR genetic elements by proportional Venn diagram (Euler; left) and bar chart (right). On right, the X axis of the bar chart depicts the presence (black points) of known AMR genes (*mphA, gyrA, parC, tet*(59), *qnr*_*Vc*_) in a given genome and the Y axis depicts the number of isolates that share the given combination of AMR genes. Coloured bars to the left indicate the number of isolate genomes encoding resistance genes to CIP (blue), AZI (pink), or DOX (yellow). AMR genetic elements to other antibiotics are not shown.

To evaluate the generalizability of these findings from the primary sample collection, we also compared aerobic and anaerobic growth curves of 277 isolates from the secondary sample collection. For ciprofloxacin, the rates of resistance were significantly different under anaerobic conditions (21%; n= 58/277) compared to aerobic conditions (1.1%; n= 3/277; McNemar’s test p<0.001; Supplementary Table 6). For azithromycin, the rates of resistance were significantly different under anaerobic conditions (100%; N= 277/277) compared to aerobic conditions (57%; N= 159/277; McNemar’s test p<0.001; Supplementary Table 6). For doxycycline, only two isolates were resistant under anaerobic conditions alone and one under both anaerobic and aerobic conditions (Supplementary Table 6). The odds of classifying isolates as resistant increased over 25 times for ciprofloxacin (OR= 25.7; 95% CI= 18.8-34.6) and 119 times for azithromycin (OR = 119; 95% CI= 20.95 – 4739) under anaerobic compared to aerobic conditions.

### Addition of catalase to test if reactive oxygen species effect antibiotic resistance/sensitivity under aerobic conditions

In this experiment, catalase was added to the media to quench hydrogen peroxide with the objective of testing the hypothesis that susceptibility under aerobic conditions was associated with ROS (e.g. hydrogen peroxide). For ciprofloxacin, the MICs for the sensitive reference strain E7946 (Cip^S^, Azi^S^, Dox^S^) and the resistant clinical isolate EN160 (Cip^R^, Azi^R^, Dox^S^) remained unchanged when catalase was added to the media under aerobic conditions. The addition of catalase was not associated with differences in AUCs for both E7946 and EN160 in media containing ciprofloxacin, azithromycin, or doxycycline at 2-fold below the MIC. The AUCs in LB media with and without catalase alone for E7946 and EN160 were not statistically different (Supplementary Table 7).

### Molecular AMR correlates under aerobic and anaerobic conditions

The distribution of known AMR genetic elements is shown (Fig. 2B). AMR-associated point mutations (likely transmitted vertically, not on an established mobilizable element), and genes on known horizontally transferred mobilizable elements, are provided (Supplementary Material). The integrative conjugative element (ICE) SXT/R391 was found in 90% (60/67) of isolates. The ICE elements contained the pentapeptide repeat protein that confers fluoroquinolone resistance (*qnr*_*Vc*_), the macrolide-inactivating phosphotransferase (*mphA*), and the major facilitator superfamily (MFS) efflux pump conferring tetracycline resistance (*tet*(59)) [52-55]. The genes *qnr*_*Vc*_, *mphA*, and *tet*(59) were found in 78% (52/66), 33% (22/66), and 78% (52/66) of isolates, respectively. Ciprofloxacin resistance under both anaerobic and aerobic conditions was significantly associated with *qnr*_*VC*_, *gyrA* and *parC* (Supplementary Table 8). Identification of the known azithromycin AMR gene *mphA* was significantly associated with resistance under aerobic conditions alone (P<0.001). The gene *tet*(59) was not associated with doxycycline resistance under aerobic or aerobic conditions (both p=0.566).

We next used GWAS to comprehensively explore the genetic basis of AMR. This approach used the phenotype of AUC to represent ‘growth’ with or without exposure to the three test antibiotics at five concentrations under either aerobic or anaerobic conditions. Growth phenotypes (analyzed by AUCs) at similar antibiotic concentrations were positively correlated within aerobic and anaerobic conditions for all three antibiotics (Fig. 3). Growth phenotypes were also positively correlated between aerobic and anaerobic conditions for ciprofloxacin (Fig. 3A). However, phenotypes were weakly, or even negatively, correlated between aerobic and anaerobic conditions for azithromycin and doxycycline (Fig. 3B,C). These results support the hypothesis that anaerobic and aerobic growth under antibiotic pressure can differ and be distinct to antibiotic type.

**FIG 3.**
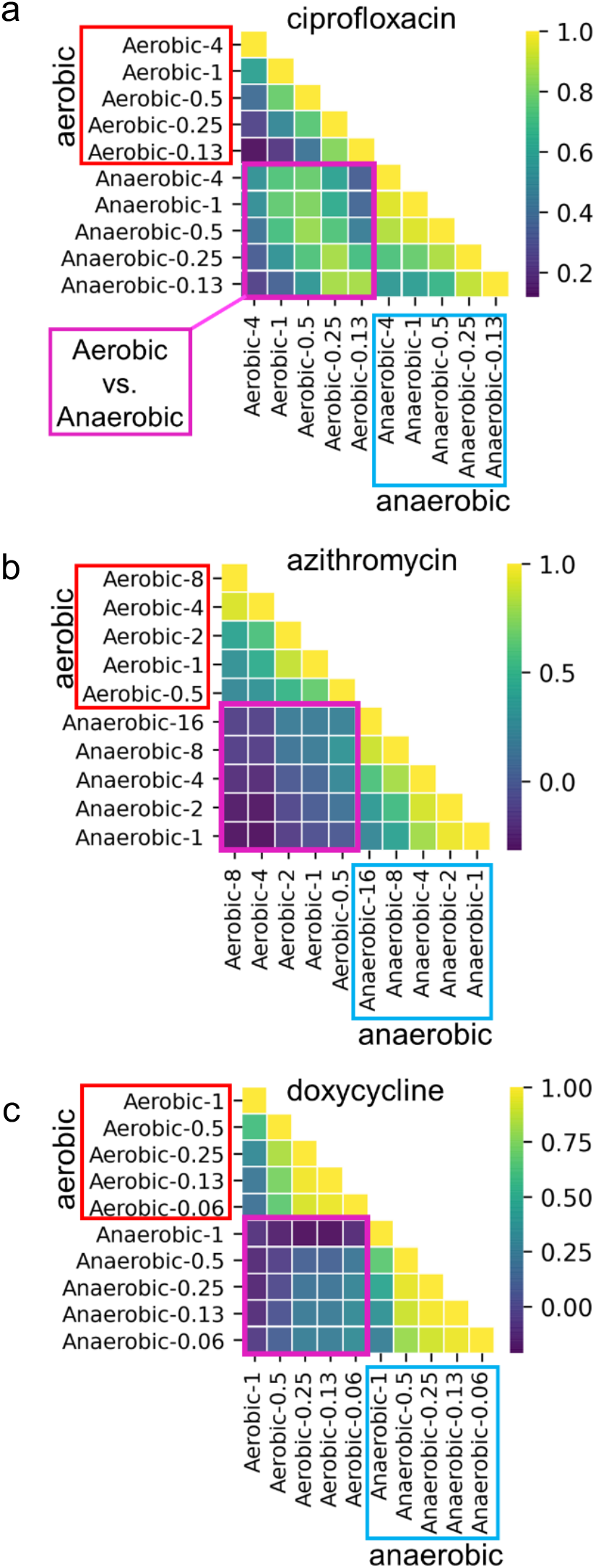
Correlation analysis of growth phenotypes at different concentrations of antibiotics under aerobic and anaerobic conditions among *V. cholerae* clinical isolates from the primary collection. Antibiotic exposures were ciprofloxacin (**a**), azithromycin (**b**), and doxycycline (**c**). AUC was analyzed as the growth parameter. Aerobic/anaerobic conditions are labeled horizontally and vertically with the antibiotic concentration in µg/ml (e.g., “Anaerobic-0.06”). Analyses are grouped: aerobic vs aerobic = red boxes; anaerobic vs aerobic = purple box; anaerobic vs anaerobic = blue boxes. Heatmaps show correlation coefficients (scale bar is to right) for similar (yellow) vs dissimilar (purple) growth at two given conditions.

The heritability of the AMR phenotypes (AUCs) was estimated prior to the GWAS. Heritability (*h*^*2*^) is defined as the proportion of phenotypic variation explained by genetic variation, measured as unique contiguous tracts of the assembled genomes (unitigs) that tag both single nucleotide variants, indels, and gene content changes (Methods). We found relatively high heritability (*h*^*2*^ in the range 0.60-0.99) of growth across concentrations of ciprofloxacin under both aerobic and anaerobic conditions, yielding statistically significant GWAS hits (Table 1; Supplementary Data Files S1 and S2): 20 under aerobic conditions and 16 under anaerobic conditions. In contrast, heritability tended to be much lower under aerobic compared to anaerobic conditions for both azithromycin and doxycycline, yielding significant GWAS hits only under anaerobic conditions (Table 1; Supplementary Data Files S3 and S4, respectively): 3 for azithromycin under anaerobic conditions alone and 57 for doxycycline under anaerobic conditions alone.

**Table 1.** Identification of genetic elements by GWAS that associate with AMR.

| Condition | Outcome <sup>b</sup> | Antibiotic Concentration (µg/ml) |  |  |  |  |
| --- | --- | --- | --- | --- | --- | --- |
| Ciprofloxacin<sup>a</sup> |  | CIP4 | CIP1 | CIP0.5 | CIP0.25 | CIP0.13 |
| Aerobic | Heritability ($h^2$) | 0.99 | 0.73 | 0.60 | 0.74 | 0.92 |
|  | Associated genes | 0 | 11 | 9 | 2 | 6 |
| Anaerobic | Heritability ($h^2$) | 0.81 | 0.73 | 0.72 | 0.72 | 0.87 |
|  | Associated genes | 8 | 8 | 8 | 10 | 4 |
| Azithromycin<sup>a</sup> |  | AZI16 | AZI8 | AZI4 | AZI2 | AZI1 |
| Aerobic | Heritability ($h^2$) | - | 0.00 | 0.02 | 0.00 | 0.11 |
|  | Associated genes | - | - | - | - | - |
| Anaerobic | Heritability ($h^2$) | 0.30 | 0.77 | 0.66 | 0.38 | 0.00 |
|  | Associated genes | 2 | 1 | 0 | 0 | - |
| Doxycycline<sup>a</sup> |  | DOX1 | DOX0.5 | DOX0.25 | DOX0.13 | DOX0.06 |
| Aerobic | Heritability ($h^2$) | 0.00 | 0.27 | 0.23 | 0.30 | 0.50 |
|  | Associated genes | - | 0 | 0 | 0 | 0 |
| Anaerobic | Heritability ($h^2$) | 0.10 | 0.61 | 0.72 | 0.72 | 0.77 |
|  | Associated genes | 0 | 23 | 55 | 54 | 17 |

<sup>a</sup> Ciprofloxacin = CIP; Azithromycin = AZI; Doxycycline = DOX
<sup>b</sup> Heritability is the proportion of phenotypic variation that is explained by genetic variation. Associated genes are all significant GWAS hits after correction for multiple hypothesis testing ( $P < 0.05$ after Bonferroni correction).

AMR genes identified by GWAS were diverse (Fig. 4; Supplementary Data Files S1-S4). These candidates included known AMR genes, such as *qnr*_*Vc*_ and *dfrA*, which were associated with ciprofloxacin resistance under both aerobic and anaerobic conditions. We identified seven genes associated with ciprofloxacin resistance under anaerobic conditions alone (including the stress response gene *barA* and a *radC* homolog involved in DNA repair; Supplementary Data File S2), and ten genes under aerobic conditions alone (including *rtxB*; Supplementary Data File S1). Under anaerobic conditions, most genes were identified at ciprofloxacin concentrations at, or above, 0.25 µg/ml; however, four genes, including *barA*, were identified under one of the lowest tested ciprofloxacin concentrations (0.13 µg/ml; Supplementary Data File S2). GWAS hits for azithromycin and doxycycline resistance were found only under anaerobic conditions. For azithromycin, two genetic elements were identified: *mphA* and a region between *ompT* and *dinG* (*ompT-dinG*; Supplementary Data File S3). For doxycycline, 23 genes were shared across concentrations; however, the gene discovery rate was highest at the lower concentrations (n=53 at 0.13 µg/ml; n =26 µg/ml; Supplementary Data File S4). GWAS hits included the major facilitator superfamily antibiotic efflux pump *tet*(59) (Fig. 4, Supplementary Data Files). Most genetic elements identified have unknown function. The identification of known AMR genes by GWAS serves as positive control and suggests that the genes of unknown function may indeed play a role in AMR.

**FIG 4.**
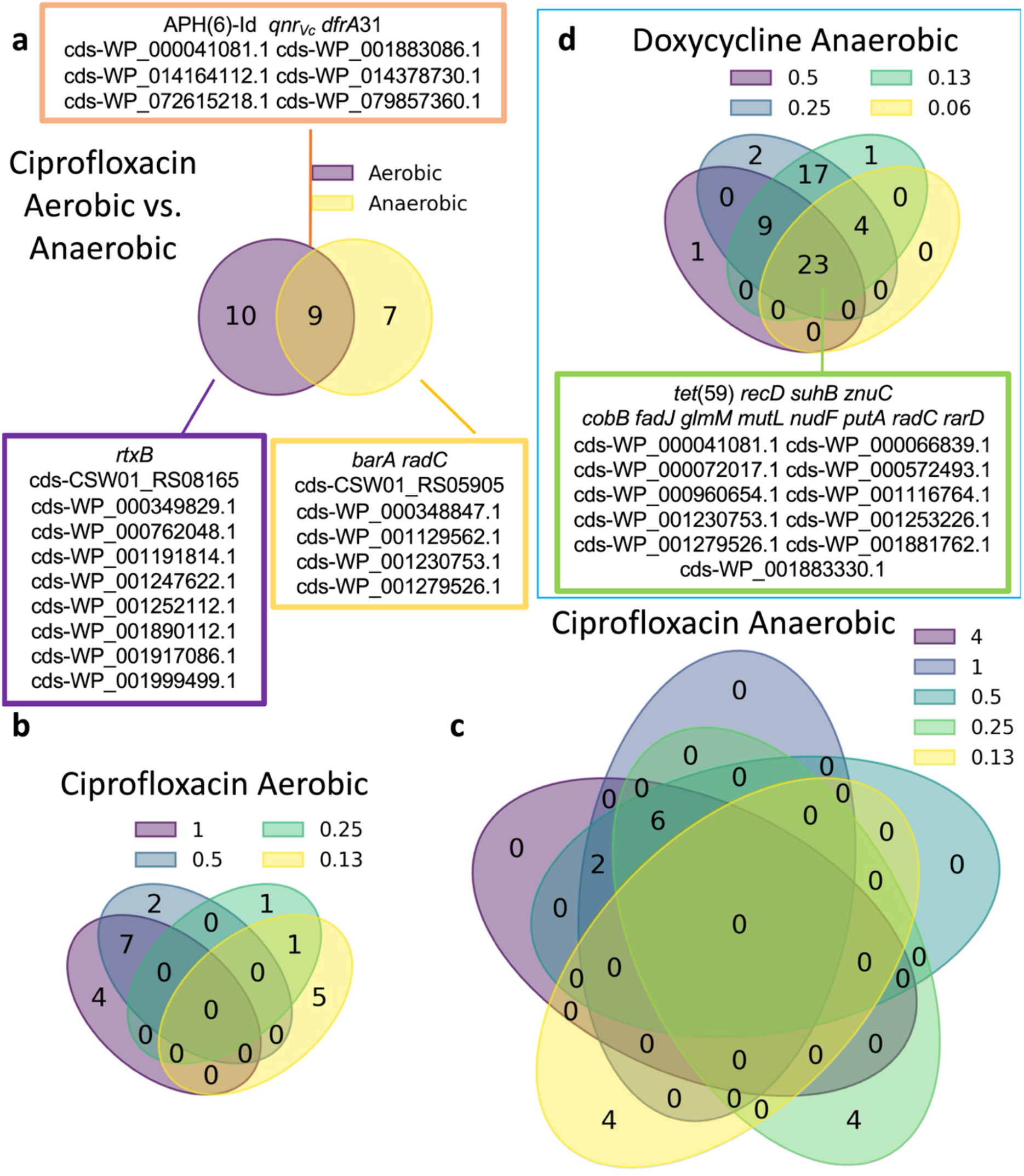
Distribution of AMR genes associated with AMR growth phenotypes at different concentrations of antibiotics under aerobic and anaerobic conditions among isolates from the primary collection. Venn diagrams show the overlap between genes associated with (**a**) ciprofloxacin resistance under aerobic vs. anaerobic conditions, (**b**) ciprofloxacin at different concentrations (µg/ml) under aerobic conditions, (**c**) ciprofloxacin at different concentrations (µg/ml) under anaerobic conditions, and (**d**) doxycycline at different concentrations (µg/ml) under anaerobic conditions. Genes shown in boxes had statistically significant associations.

### Antibiotics detected in stool by LC-MS/MS

Finally, we sought to test the hypothesis that AMR genotypes and phenotypes would be associated with measured concentrations of antibiotics in stool. A combined total of 196 antibiotics were detected in the 51 stool supernatants tested by mass spectrometry using a targeted technique for 9 common antibiotics (Fig. 5). At least one antibiotic was detected in 98% (n=50/51), at least two antibiotics were detected in 94% (n=48/51), and three or more antibiotics were detected in 90% (n=46/51) of stool supernatants (Fig. 5). Antibiotics detected were ciprofloxacin (n=48/51; 94%), tetracycline / doxycycline (n=46/51; 90%), nalidixic acid (n=41/51; 80%), metronidazole (n=37/51; 73%), sulfamethoxazole/trimethoprim (n=22/51; 43%), and amoxicillin (n=2/51; 4%); azithromycin was not detected. Detection of quinolone/fluoroquinolone and tetracycline/doxycycline in stool by LC-MS/MS was not associated with AMR genotypes or phenotypes (Supplementary Table 9). Associations for azithromycin could not be tested because azithromycin was not detected in any stool supernatant.

**FIG 5.**
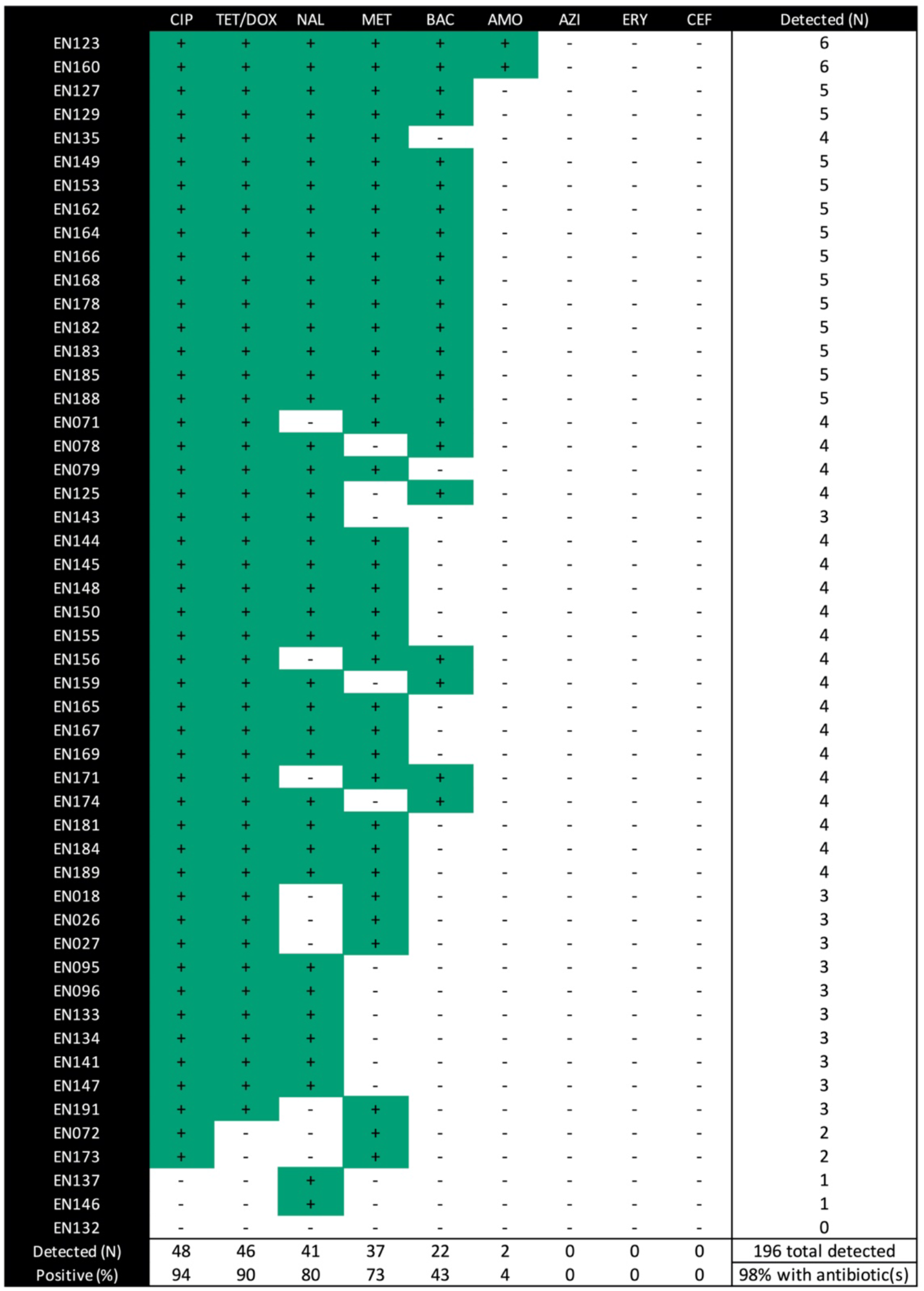
Antibiotic detection in stool supernatants by mass spectrometry (LC-MS/MS) among cholera samples from the primary collection. Green with “+” = Detected. White with “-” = not detected. CIP= ciprofloxacin, TET/DOX= tetracycline and/or doxycycline, NAL = nalidixic acid, MET = metronidazole, BAC = sulfamethoxazole and/or trimethoprim, AMO = amoxicillin, ERY = erythromycin, CEF = ceftriaxone. Stool supernatants were not available for EN80, 86, 88, 92, 100, 103, 109, 116-120, 126, 124, 130, 131.

## DISCUSSION

In this study, *V. cholerae* isolates from cholera patients were found to be more resistant to antibiotic exposure under anaerobic conditions compared to aerobic conditions (Fig. 1). This phenotype differed by antibiotic class. Novel genetic elements were found to associate with AMR under anaerobic conditions which also differed by antibiotic class (Fig. 4). The approach of using a continuous variable (area under the growth curve (AUC)) for the AMR phenotype within the framework of GWAS may provide a new approach to identify putative AMR genetic targets for future mechanistic research.

CLSI breakpoints for enteropathogens like *V. cholerae* were developed under aerobic conditions [45, 56]. CLSI and other clinical reference bodies set breakpoints to have clinical relevance despite limited data from clinical studies [56-58]. In this context, the odds of classifying isolates in the primary collection as resistant under anaerobic conditions compared to aerobic conditions increased over 10 times for ciprofloxacin and over 200 times for azithromycin. These results are likely general across *V. cholerae* because in the secondary collection, which is separated by more than 10 years, we found that the odds of classifying isolates as resistant under anaerobic conditions compared to aerobic conditions increased over 20 times for ciprofloxacin and over 100 times for azithromycin.

There are several physiologic explanations for increased antibiotic resistance under anaerobic conditions. ROS induce both intracellular and cell-wall stress and are at higher concentrations under aerobic conditions [59]. ROS may have acted synergistically to potentiate antibiotic lethality [31, 60]. The assays that utilized catalase to quench hydrogen peroxide under aerobic conditions were conducted to evaluate this possibility. The MICs for all three antibiotics did not increase with the addition of catalase suggesting that the reduction of ROS alone cannot account for increased MICs observed under anaerobic conditions (Supplementary Table 7). We hypothesize that reduced growth under anaerobic conditions might decrease the effectiveness of antimicrobial agents that directly or indirectly disrupt cell envelope integrity [61]. While reduced growth was observed under anaerobic conditions, the antibiotics tested are not known to directly disrupt the cell envelope. However, off target effects (e.g. cell envelope stress) by CIP, AZI, and DOX may have occurred.

To further investigate AMR phenotypes under anaerobic conditions, future studies will benefit from the use of a defined medium (such as M9) where the carbon source and an alternative electron acceptor (e.g. fumarate, nitrate, dimethylsulfoxide, or trimethylamine N-oxide) can be supplemented to strictly control anaerobic respiration versus fermentation. Buffering the medium (e.g. with PBS or bicarbonate) to a neutral pH is important because *V. cholerae* can acidify the medium over time, with toxic effects. In prior research, acidification occurred after 10 hours [9]. In our shorter 8-hour assay, acidification was observed under anaerobic conditions among 1 of 4 strains tested (Supplementary Table 3). Furthermore, buffering to the alkalinic pH found in cholera stool (pH 8.5-9) will provide important insight given that *V. cholerae* utilizes nitrate for anaerobic respiration only at alkaline pH [2, 9]. Here we used an undefined medium (LB) without a defined, saturating concentration of an alternative electron acceptor. This approach likely resulted in mixed anaerobic respiration and fermentation. To address these limitations in future studies, robotic automation and 384-well formatted assays would enable a scalable system for multiple defined media across broad gradients of antibiotics. Ideally, we would simulate the conditions of the human gut, but in practice these conditions can only be approximated.

There are many knowledge gaps on the genetic basis of the AMR phenotypes under varying environmental conditions. This study prioritized the factor of oxygen exposure as a determinant of AMR phenotypes because facultative anaerobic enteropathogens experience hypoxia and anoxia in the animal gut [9]. The first phase of the analysis focused on previously known AMR genotypes with known AMR phenotypes. For ciprofloxacin, mutations in *parC* and carriage of *qnr*_*Vc*_ were significantly associated with phenotypic resistance under aerobic and anaerobic conditions; mutations in *gyrA* were significantly associated with resistance under anaerobic conditions alone (Supplementary Table 8). For azithromycin, *mphA* was identified and significantly associated with AMR under aerobic conditions alone. While *tet*(59) was identified, very few isolates were identified as resistant to doxycycline under aerobic (n=1) or anaerobic conditions (n=2). These associative data begin to reveal a difference between AMR genotypes and phenotypes under aerobic and anaerobic conditions.

The second phase of analysis sought to use GWAS to identify previously unknown genetic targets associated with AMR. The continuous variable of AUC, as opposed to the binary variable of growth/ no growth, was used in assays with and without antibiotic exposure under aerobic and anaerobic conditions. Antibiotics at the breakpoint and sub-breakpoint concentrations were chosen based on a rationale that different genetic elements might contribute differently to AMR phenotypes at different antibiotic concentrations. As expected, GWAS identified *qnr*_*Vc*_ for ciprofloxacin exposure under both aerobic and anaerobic conditions (Fig. 4). GWAS identified *mphA* for azithromycin exposure under anaerobic conditions alone and *tet*(59) for doxycycline exposure under anaerobic conditions alone. These results of known AMR genes served as ‘positive controls’ for the GWAS. We also note that genes known to be important for anaerobic respiration (e.g. *tatA1, tatC, ccmA-F, ccmH, moaA-D, moeA-B, napA-D, napF, fnr, narP-Q, nqrF, dsbA, dsbD, hemN*) in *V. cholerae* were not identified as GWAS hits [9]. This suggests that our GWAS was specific to AMR phenotypes and was not liable to detect genes related to anaerobic conditions alone. We therefore expect novel GWAS hits to be likely candidates for involvement in AMR phenotypes.

For ciprofloxacin, seven genes were associated with AMR in anaerobic conditions alone; these included a gene involved in DNA repair (*radC*), 2-component histidine kinase involved in stress response (*barA*), and an ATP-dependent zinc protease. The gene *dfrA31* encodes a trimethoprim-resistant dihydrofolate reductase and APH(6)-Id encodes a streptomycin phosphotransferase enzyme; both genes were identified for aerobic and anaerobic conditions. These two genes are located on the SXT element along with *qnr*_*Vc*_ and may therefore be associated due to genetic linkage rather than due to causal roles in ciprofloxacin resistance. For azithromycin, one additional genetic element under anaerobic conditions was discovered to associate with AMR: an intergenic region between *ompT* (porin; known to be associated with AMR) [62, 63] and *dinG* (ATP-dependent DNA helicase). For doxycycline, a diverse set of 57 genetic elements under anaerobic conditions alone were discovered to associate with AMR. These include *vexK* (efflux RND transporter permease associated with AMR) [64-66], and *zorA* (anti-phage defense system ZorAB subunit A; a putative proton channel that may respond to membrane perturbation by depolarization) [67].

In addition to the SXT element, genes associated with an AMR phenotype were also discovered on the Vibrio Pathogenicity Island II (VSPII; N16961 VC0506-VC0512 / E7946 loci RS02705-RS02745); these loci are genetically diverse in Bangladesh [68]. The GWAS hits in VSPII encode both biofilm/ auto-aggregation associated factors as well as an aerotaxis protein (AerB; VC0512) [69]; findings consistent with roles in AMR and aerobic/anaerobic conditions. These GWAS analyses were of limited power due to the modest sample size, and could be sensitive to false positives at AMR ‘hot-spots’ like SXT. Despite these limitations, GWAS enabled the discovery of an intriguing list of genetic targets that were associated with AMR and require future mechanistic molecular analysis to test for causal relationships.

LC-MS/MS analysis on the stools from the primary collection, stools from which the isolates were obtained, was conducted to test the hypothesis that the rates of AMR genotypes and phenotypes were higher when the stool samples contained the cognate antibiotic. Nearly all patients shed at least one antibiotic, making it difficult to identify AMR correlates to exposure (Fig. 5). This finding is important because studies that leverage natural infection to set clinically meaningful AMR breakpoints under aerobic conditions, and now anaerobic conditions, cannot readily be performed because of the degree of antibiotic exposure among diarrheal patients. Therefore, future interventional clinical studies with known antibiotic exposure determined *a priori* may be required. Given that the primary collection is from patients that self-reported not taking antibiotics, the detection of a combined total of 196 antibiotics further highlights the ubiquity of antibiotics and the limited value of self-reported antibiotic exposure.

## Conclusions

Facultative enteropathogens are exposed to antibiotics under aerobic and anaerobic conditions in both the human gut and in the environment. We used the facultative anaerobic enteropathogen *V. cholerae* as a model to test for differences in AMR phenotypes under aerobic and anaerobic conditions. Increased resistance was found under anaerobic conditions compared to aerobic conditions. Using AMR breakpoints established for aerobic conditions, the odds of classifying isolates as resistant under anaerobic compared to aerobic conditions increased over 10 times for two of the three antibiotics tested. While several known resistance genes were associated with AMR under both conditions, many genes were only associated with AMR under one condition. Heritability tended to be higher, and more genes associated with resistance, under anaerobic conditions. This suggests that key genetic determinants of resistance may be missed when experiments are only performed aerobically. Our findings provide a rationale to determine if increased MICs under anaerobic conditions are associated with therapeutic failures and/or microbial escape in cholera patients, and if true, there may be a need to determine AMR breakpoints for anaerobic conditions.

## Acknowledgements

We thank the patients for participating in this study and the icddr,b clinical and laboratory teams that collected the samples. We are grateful to Randy Autrey and Krista Berquist for their administrative expertise, as well as Glenn Morris at the Emerging Pathogens Institute and Desmond Schatz in the Department of Pediatrics at the University of Florida for their ongoing support. Stephen Calderwood, Jason Harris, and Regina LaRocque were the principal investigators (PI) of the parent study/IRB protocol (Massachusetts General Hospital, Harvard University School of Medicine, USA) under which the samples were collected by EJN when he was a NIH Fogarty Fellow. Andrew Camilli provided additional reagents and insights for this manuscript; FQ was the principal investigator in Bangladesh and PI of the ERC/RRC approvals at the icddr,b. This generous research infrastructure and support was invaluable to the success of this study.

## Disclaimer

The funders had no role in study design, data collection and analysis, decision to publish, or preparation of the manuscript.

