## Supplemental Material for "Genome-wide association studies reveal distinct genetic correlates and increased heritability of antimicrobial resistance in *Vibrio cholerae* under anaerobic conditions"

**SUPPLEMENTARY MATERIAL**

**Supplementary Data Files (excel file with four datafile tabs)**

Date File S1. GWAS Hits. Ciprofloxacin resistance phenotype. Aerobic.
Date File S2. GWAS Hits. Ciprofloxacin resistance phenotype. Anaerobic.
Data File S3. GWAS Hits. Azithromycin resistance phenotype. Anaerobic.
Data File S4. GWAS Hits. Doxycycline resistance phenotype. Anaerobic.

**Supplementary Results.**

**Antibiotic resistance phenotypes of known AMR genetic elements under aerobic and anaerobic conditions**. The distribution of known AMR genetic elements (Figure 2) was grouped by point mutations (likely transmitted vertically, not on an established mobilizable element) and horizontal transmission (on an established mobilizable element).

*Point mutations in known AMR genes.* For ciprofloxacin, the most identified known resistance mutations were those in the topoisomerase encoding genes *gyrA* (VC1258/RS06370) (Ser83Ile) and *parC* (VC2430/RS12340) (Ser85Leu). These were present in 60% (40/67) and 70% (47/67) of all isolates, while 54% (36/57) contained both. For azithromycin, we found no mutations in known resistance genes encoding ribosomal proteins L4 (*rplD;*VC2595/RS13175) and L22 (*rplV;*VC2591/RS13155) [1, 2]. For tetracycline (proxy for doxycycline), one of the 9 total 16S rRNA genes (VCr001) was found to have a single nucleotide insertion of a G nucleotide at position 327 within the sequences of 15% (10/67) of isolates; the significance is unknown. No mutations were detected in the 30S ribosomal protein encoded by genes *rpsJ* (VC2597/RS13185) or *rpsC* (VC2590/RS13150), whose mutations have been associated with tetracycline class resistance in other Gram negative organisms [3-5].

*Known AMR genes on mobile genetic elements.* The integrative conjugative element (ICE) SXT/R391 was found in 90% (60/67) of isolates. The ICE contained a pentapeptide repeat protein conferring fluoroquinolone resistance ($qnr_{Vc}$), a macrolide-inactivating phosphotransferase (*mphA*), and a major facilitator superfamily (MFS) efflux pump conferring tetracycline resistance (*tet*(59)) [6-9]. The genes $qnr_{Vc}$, *mphA*, and *tet*(59) were also found in 78% (52/67), 33% (22/67), and 78% (52/67) of isolates, respectively.

**Supplementary Table 1.** Reference strains and clinical isolates

| | | Aerobic[b] | | | Anaerobic[b] | | | Source[c] |
|---|---|---|---|---|---|---|---|---|
| | Strain[a] | $CIP^R$ | $AZI^R$ | $DOX^R$ | $CIP^R$ | $AZI^R$ | $DOX^R$ | |
| Reference | E7946 | - | - | - | - | + | - | (1) |
| Clinical Isolates | | | | | | | | |
| EN018 | L_EN1286 | - | - | - | - | + | - | (2) |
| EN026 | L_EN1291 | - | - | - | - | + | - | (2) |
| EN027 | L_EN1292 | - | - | - | - | + | - | (2) |
| EN071 | L_EN1300 | + | - | - | + | + | - | (2) |
| EN072 | L_EN1301 | - | - | - | + | + | - | (2) |
| EN078 | L_EN1303 | + | - | - | + | + | - | (2) |
| EN079 | L_EN1304 | - | + | - | + | + | - | (2) |
| EN080 | L_EN1305 | + | - | - | + | + | - | (2) |
| EN086 | L_EN1307 | + | + | - | + | + | - | (2) |
| EN088 | L_EN1308 | + | - | - | + | + | - | (2) |
| EN092 | L_EN1310 | + | - | - | + | + | - | (2) |
| EN095 | L_EN1312 | + | + | - | + | + | - | (2) |
| EN096 | L_EN1313 | + | + | - | + | + | - | (2) |
| EN100 | L_EN1314 | + | + | - | + | + | - | (2) |
| EN103 | L_En1315 | - | - | - | + | + | - | (2) |
| EN109 | L_EN1319 | - | - | - | + | + | - | (2) |
| EN116 | L_EN1325 | - | - | - | + | + | - | (2) |
| EN117 | L_EN1326 | - | - | - | - | + | - | (2) |
| EN118 | L_EN1327 | - | + | - | + | + | - | (2) |
| EN119 | L_EN1328 | + | - | - | + | + | - | (2) |
| EN120 | L_EN1329 | - | - | - | + | + | - | (2) |
| EN123 | L_EN1330 | - | - | - | + | + | - | (2) |
| EN124 | L_EN1331 | + | - | - | + | + | - | (2) |
| EN125 | L_EN1332 | + | + | - | + | + | - | (2) |
| EN126 | L_EN1333 | + | - | - | + | + | - | (2) |
| EN127 | L_EN1334 | + | - | - | + | + | - | (2) |
| EN129 | L_EN1335 | - | - | - | + | + | - | (2) |
| EN130 | L_EN1336 | + | - | - | + | + | - | (2) |
| EN131 | L_EN1337 | + | - | - | + | + | - | (2) |
| EN132 | L_EN1338 | + | - | - | + | + | - | (2) |
| EN133 | L_EN1339 | + | - | - | + | + | - | (2) |
| EN134 | L_EN1340 | - | + | - | + | + | - | (2) |

| | | | | | | | | |
|---|---|---|---|---|---|---|---|---|
| EN135 | L_EN1341 | - | - | - | + | + | - | (2) |
| EN137 | L_EN1343 | + | + | - | + | + | - | (2) |
| EN141 | L_EN1344 | + | - | - | + | + | - | (2) |
| EN143 | L_EN1346 | - | - | - | + | + | - | (2) |
| EN144 | L_EN1347 | + | - | - | + | + | - | (2) |
| EN145 | L_EN1348 | + | + | - | + | + | + | (2) |
| EN146 | L_EN1349 | + | - | - | + | + | - | (2) |
| EN147 | L_EN1350 | - | - | - | + | + | - | (2) |
| EN148 | L_EN1351 | - | - | - | + | + | - | (2) |
| EN149 | L_EN1352 | + | - | - | + | + | - | (2) |
| EN150 | L_EN1353 | - | - | - | + | + | - | (2) |
| EN153 | L_EN1355 | - | - | - | + | + | - | (2) |
| EN155 | L_EN1357 | + | - | - | + | + | - | (2) |
| EN156 | L_EN1358 | - | - | - | + | + | - | (2) |
| EN159 | L_EN1360 | + | + | - | + | + | - | (2) |
| EN160 | L_EN1361 | + | - | - | + | + | - | (2) |
| EN162 | L_EN1363 | + | + | - | + | + | - | (2) |
| EN164 | L_EN1365 | - | - | - | + | + | - | (2) |
| EN165 | L_EN1366 | + | - | - | + | + | - | (2) |
| EN166 | L_EN1367 | + | - | - | + | + | - | (2) |
| EN167 | L_EN1368 | + | - | - | + | + | - | (2) |
| EN168 | L_EN1369 | + | + | - | + | + | - | (2) |
| EN169 | L_EN1370 | - | - | - | + | + | - | (2) |
| EN171 | L_EN1371 | + | + | + | + | + | + | (2) |
| EN173 | L_EN1372 | - | - | - | + | + | - | (2) |
| EN174 | L_EN1374 | - | - | - | + | + | - | (2) |
| EN178 | L_EN1377 | + | - | - | + | + | - | (2) |
| EN181 | L_EN1379 | - | - | - | - | + | - | (2) |
| EN182 | L_En1380 | - | - | - | + | + | - | (2) |
| EN183 | L_EN1381 | - | - | - | + | + | - | (2) |
| EN184 | L_EN1382 | + | - | - | + | + | - | (2) |
| EN185 | L_EN1383 | - | - | - | + | + | - | (2) |
| EN188 | L_EN1385 | - | - | - | + | + | - | (2) |
| EN189 | L_EN1386 | + | + | - | + | + | - | (2) |
| EN191 | L_EN1388 | - | - | - | + | + | - | (2) |

[a] Prefix of "L_" was used to distinguish the strain number in the library from the clinical isolate number, which is maintained to be consistent with prior publications.

[b] $\text{CIP}^{R}$ = ciprofloxacin resistance. $\text{AZI}^{R}$ = azithromycin resistance. $\text{DOX}^{R}$ = doxycycline resistance.

[c] Sources of strains: (1) Mekalanos, J. J. Duplication and amplification of toxin genes in Vibrio cholerae. Cell 35, 253-263, (1983) ; (2) Nelson, E. J. et al. Complexity of rice-water stool from patients with Vibrio cholerae plays a role in the transmission of infectious diarrhea. Proc Natl Acad Sci U S A 104, 19091-19096, (2007).

**Supplementary Table 2.** Baseline growth parameters of *V. cholerae* clinical isolates under aerobic and anaerobic conditions

| Growth parameter [a] | Aerobic median | Anaerobic median | P [b] |
|---|---|---|---|
| K | 1.08 | 0.261 | **< 0.001** |
| AUC | 5.15 | 1.58 | **< 0.001** |
| Velocity | 0.011 | 0.006 | **< 0.001** |

[a] AUC = area under the curve (optical density x time in minutes). K = carrying capacity. Media was LB alone without antibiotics. Velocity = growth rate in percent increase per minute at half the carrying capacity.

[b] Wilcoxon signed-rank test for growth. Bold = statistically significant ($P<0.05$).

**Supplementary Table 3.** Growth and pH metrics with and without 20mM fumarate supplementation.

| | | Anaerobic | | | | Aerobic | | | |
|---|---|---|---|---|---|---|---|---|---|
| | | Replicate 1 | | Replicate 2 | | Replicate 1 | | Replicate 2 | |
| Strain | Fumarate | AUC[a] | pH[b] | AUC[a] | pH[b] | AUC[a] | pH[c] | AUC[a] | pH[c] |
| **E7946** | **-** | 1.63 | 7.2 | 1.53 | 7.2 | 5.11 | 7.2 | 5.42 | 7.2 |
| | **+** | 1.93 | 7.2 | 1.85 | 7.2 | 5.33 | 7.2 | 5.52 | 7.2 |
| **EN145** | **-** | 1.28 | 6.4 | 1.22 | 6.4 | 4.96 | 7.2 | 5.10 | 7.2 |
| | **+** | 1.65 | 7.2 | 1.55 | 7.2 | 4.98 | 7.2 | 5.33 | 7.2 |
| **EN160** | **-** | 1.55 | 6.8 | 1.33 | 6.8 | 4.63 | 7.2 | 5.03 | 7.2 |
| | **+** | 1.92 | 7.2 | 1.66 | 7.2 | 5.03 | 7.2 | 5.23 | 7.2 |
| **EN181** | **-** | 1.25 | **6.0** | 1.20 | **6.0** | 5.02 | 7.2 | 5.10 | 7.2 |
| | **+** | 1.65 | 6.8 | 1.63 | 7.2 | 5.38 | 7.2 | 5.30 | 7.2 |
| Average (st.dev) | **-** | 1.43 (0.19) | … | 1.32 (0.15) | … | 4.93 (0.21) | … | 5.16 (0.17) | … |
| | **+** | 1.79 (0.16) | … | 1.67 (0.12) | … | 5.18 (0.21) | … | 5.35 (0.12) | … |
| Percent difference with fumarate | | +25% | … | +27% | … | +5.1% | … | +3.7% | … |

[a] 'AUC' = area under the curve (optical density x time in minutes).

[b] Bold numbers represents pH of test wells below the pH of the LB blank control well at the end of the assay (pH 6.4); the pH of the LB blank control well at the start of the assay was 6.8.

[c] The starting and ending pH of the LB control blank wells were 6.8 and 7.2 for both replicates, respectively.

**Supplementary Table 4.** Minimal inhibitory concentrations (MICs) for ciprofloxacin, azithromycin, and doxycycline among *V. cholerae* clinical isolates

| | Ciprofloxacin[b] | | Azithromycin[b] | | Doxycycline[b] | |
|---|---|---|---|---|---|---|
| μg/ml[a] | Aerobic | Anaerobic | Aerobic | Anaerobic | Aerobic | Anaerobic |
| 0.002 | 0 | 0 | 0 | 0 | 0 | 0 |
| 0.004 | 1 | 0 | 0 | 0 | 0 | 0 |
| 0.008 | 0 | 0 | 0 | 0 | 0 | 0 |
| 0.016 | 2 | 1 | 0 | 0 | 0 | 0 |
| 0.032 | 0 | 0 | 0 | 0 | 0 | 0 |
| 0.063 | 0 | 1 | 0 | 0 | 0 | 0 |
| 0.13 | 0 | 1 | 0 | 0 | 2 | 1 |
| 0.25 | 0 | 0 | 0 | 0 | 7 | 4 |
| 0.5 | 5 | 0 | 0 | 0 | 14 | 9 |
| 1 | 23 | 2 | 5 | 0 | **38** | **26** |
| 2 | **35** | 4 | 22 | 0 | 2 | 21 |
| 4 | 0 | 1 | **23** | 0 | 3 | 4 |
| 8 | 1 | **56** | 15 | 3 | 1 | 1 |
| 16 | 0 | 0 | 0 | 4 | 0 | 1 |
| 32 | 0 | 1 | 2 | **44** | 0 | 0 |
| 64 | 0 | 0 | 0 | 14 | 0 | 0 |
| 124 | 0 | 0 | 0 | 2 | 0 | 0 |

[a] Concentration of antibiotic.

[b] Distribution of MICs for clinical isolates grown under aerobic and anaerobic conditions. Bold text signifies the concentration at which the MIC mode was determined among the clinical isolates.

**Supplementary Table 5.** Comparison of rates of resistance detected under aerobic versus anaerobic conditions among *V. cholerae* clinical isolates in the primary collection

| | N | $R^{Ae}/R^{An}$ | $R^{Ae}/S^{An}$ | $S^{Ae}/R^{An}$ | $S^{Ae}/S^{An}$ | p[b] |
|---|---|---|---|---|---|---|
| Ciprofloxacin[a] | 67 | 36 | 0 | 26 | 5 | **<0.001** |
| Azithromycin[a] | 67 | 15 | 0 | 52 | 0 | **<0.001** |
| Doxycycline[a] | 67 | 1 | 0 | 1 | 65 | 1 |

[a] Distribution of paired resistant ('R') and sensitive ('S') phenotypes for isolates under aerobic ('Ae') and anaerobic ('An') conditions. R is defined as an MIC at or above the CLSI breakpoints (see methods).
[b] McNemar's Exact Test. Bold = statistically significant (P<0.05)

**Supplementary Table 6.** Comparison of rates of resistance detected under aerobic versus anaerobic conditions among *V. cholerae* clinical isolates in the secondary collection

| | N | $R^{Ae}/R^{An}$ | $R^{Ae}/S^{An}$ | $S^{Ae}/R^{An}$ | $S^{Ae}/S^{An}$ | p[b] |
|---|---|---|---|---|---|---|
| Ciprofloxacin[a] | 277 | 2 | 1 | 56 | 218 | **<0.001** |
| Azithromycin[a] | 277 | 159 | 0 | 118 | 0 | **<0.001** |
| Doxycycline[a] | 277 | 1 | 0 | 3 | 273 | 0.371 |

[a] Distribution of paired resistant ('R') and sensitive ('S') phenotypes for isolates under aerobic ('Ae') and anaerobic ('An') conditions. R is defined as an MIC at or above the CLSI breakpoints (see methods).
[b] McNemar's Exact Test. Bold = statistically significant (P<0.05)

**Supplementary Table 7.** Effect of catalase on growth parameters for *V. cholerae* E7946 and EN160 under aerobic conditions

| Experiment | Strain[a] | Antibiotic[b] | Catalase[c] | AUC mean[d] | AUC IQR[d] | P[e] |
|---|---|---|---|---|---|---|
| 1. | E7946 | CIP | YES | 0.697 | 0.029 | 0.136 |
| | | CIP | NO | 0.736 | 0.014 | |
| 2. | E7946 | AZI | YES | 3.26 | 0.056 | 0.533 |
| | | AZI | NO | 3.29 | 0.005 | |
| 3. | E7946 | DOX | YES | 5.14 | 0.105 | 0.959 |
| | | DOX | NO | 5.14 | 0.070 | |
| 4. | E7946 | NO | YES | 5.29 | 0.078 | 0.818 |
| | | NO | NO | 5.30 | 0.090 | |
| 5. | EN160 | CIP | YES | 2.31 | 0.368 | 0.551 |
| | | CIP | NO | 2.10 | 0.420 | |
| 6. | EN160 | AZI | YES | 4.43 | 0.119 | 0.571 |
| | | AZI | NO | 4.50 | 0.113 | |
| 7. | EN160 | DOX | YES | 4.67 | 0.322 | 0.895 |
| | | DOX | NO | 4.60 | 0.360 | |
| 8 | EN160 | NO | YES | 5.33 | 0.188 | 0.092 |
| | | NO | NO | 5.44 | 0.146 | |

[a] E7946 ($Cip^S$, $Azi^S$,$Dox^S$) is the reference strain and EN160 ($Cip^R$, $Azi^R$, $Dox^S$) is a clinical isolate. Biological replicates in experiments 1-3 and 5-7 were 3, each with 4 technical replicates. Biological replicates for experiments 4 and 8 were 9, each with 4 technical replicates.

[b] CIP = ciprofloxacin. AZI = azithromycin. DOX = doxycycline. Assays were run at CIP = 0.5, AZI = 2, and DOX = 0.25 µg/ml for EN160; E7946 was run at CIP = 0.002, AZI = 1, and DOX = 0.013 µg/ml.

[c] CIP and AZI were tested with 3 biological replicates; DOX with 2 biological replications; LB controls with 9 biological replicates.

[d] AUC = area under the curve. IQR = interquartile range.

[e] Student's t-test. Bold = statistically significant ($P<0.05$)

**Supplementary Table 8.** Comparison of antibiotic resistance phenotypes and known resistance genotypes among *V. cholerae* clinical isolates in the primary collection

Aerobic conditions [a]

| Antibiotic tested | Gene | R/P | R/NP | S/P | S/NP | OR[b] | 95% CI[b] | P[b] |
|---|---|---|---|---|---|---|---|---|
| Ciprofloxacin | *qnr*$_{Vc}$ | 36 | 0 | 16 | 15 | 31 | (4.22 - 695) | **<0.001** |
| | *gyrA* | 28 | 8 | 12 | 19 | 5.4 | (1.86 - 17.9) | **0.002** |
| | *parC* | 31 | 5 | 16 | 15 | 5.6 | (1.65 - 18.9) | **0.003** |
| Azithromycin | *mphA* | 15 | 0 | 7 | 45 | 83 | (11.2 - 1938) | **<0.001** |
| Doxycycline | *tet*(59) | 1 | 0 | 51 | 15 | 0.62 | (0.046 - 18.9) | 0.566 |

Anaerobic conditions [a]

| Antibiotic tested | Gene | R/P | R/NP | S/P | S/NP | OR[b] | 95% Cb[c] | P[b] |
|---|---|---|---|---|---|---|---|---|
| Ciprofloxacin | *qnr*$_{Vc}$ | 52 | 10 | 0 | 5 | 27 | (3.53 - 666) | **<0.001** |
| | *gyrA* | 40 | 22 | 0 | 5 | 10 | (1.39 - 248) | **0.016** |
| | *parC* | 47 | 15 | 0 | 5 | 17 | (2.28 - 416) | **0.003** |
| Azithromycin | *mphA* | 22 | 45 | 0 | 0 | 0.51 | (0.0127 - 20.1) | 1 |
| Doxycycline | *tet*(59) | 1 | 0 | 51 | 15 | 0.62 | (0.046 - 18.9) | 0.566 |

[a] Whole genome sequencing data from the primary collection were analyzed for known AMR genes. R = resistant phenotype, S = sensitive phenotype, P = gene present, NP = gene not present. Enumerations include isolates with at least the specific gene named; isolates may have more than one resistance gene (e.g. *qnr*$_{Vc}$, *gyrA* and *parC*).

[b] Fisher's Exact Test. Bold = statistically significant (P<0.05)

**Supplementary Table 9.** Test of association between antibiotic detection by mass spectrometry and AMR genotypes and phenotypes among *V. cholerae* clinical isolates

Antibiotic detection (D) and AMR genotype present (P) [a]

| | D/P | D/NP | ND/P | ND/NP | *P*[b] |
|---|---|---|---|---|---|
| CIP | 41 | 7 | 3 | 0 | 1 |
| CIP + NAL | 35 | 4 | 9 | 3 | 0.334 |
| DOX | 16 | 8 | 20 | 7 | 0.759 |
| DOX + TET | 2 | 0 | 34 | 15 | 1 |

Antibiotic detection (D) and resistance phenotype (R) under aerobic conditions [a]

| | D/R | D/S | ND/R | ND/S | *P*[b] |
|---|---|---|---|---|---|
| CIP | 24 | 24 | 3 | 0 | 0.238 |
| CIP + NAL | 22 | 17 | 5 | 7 | 0.511 |
| DOX | 1 | 23 | 0 | 27 | 0.471 |
| DOX + TET | 0 | 2 | 1 | 48 | 1 |

Antibiotic detection (D) and resistance phenotype (R) under anaerobic conditions [a]

| | D/R | D/S | ND/R | ND/S | *P*[b] |
|---|---|---|---|---|---|
| CIP | 44 | 4 | 3 | 0 | 1 |
| CIP + NAL | 38 | 1 | 9 | 3 | **0.036** |
| DOX | 1 | 23 | 0 | 27 | 0.217 |
| DOX + TET | 0 | 2 | 1 | 48 | 1 |

[a] D=detected, ND=not detected, R=resistant by MIC, S=sensitive by MIC, P = Present, NP = Not present.

[b] Fisher's Exact Test. Bold = statistically significant (P<0.05)
